# Copy number variants in the sheep genome detected using multiple approaches

**DOI:** 10.1101/041475

**Authors:** Gemma M Jenkins, Michael E Goddard, Michael A Black, Rudiger Brauning, Benoit Auvray, Ken G Dodds, James W Kijas, Noelle Cockett, John C McEwan

**Affiliations:** AbacusBio Limited, 442 Moray Place, PO Box 5585, Dunedin 9058, NEW ZEALAND; Victorian Department of Economic Development, Jobs, Transport and Resources, Bundoora, VIC 3083, AUSTRALIA; Department of Biochemistry, University of Otago, 710 Cumberland St, Dunedin 9054, NEW ZEALAND; AgResearch, Invermay Agricultural Centre, PB 50034, Mosgiel 9053, NEW ZEALAND; CSIRO Animal, Food and Health Sciences, Queensland Bioscience Precinct, 306 Carmody Road. St Lucia, QLD 4067, AUSTRALIA; Utah State University, 1435 Old Main Hill, Logan, UT 84322-1435-1435, USA

**Keywords:** sheep, copy number variants, array CGH, SNP, sequence

## Abstract

**Background.:** Copy number variants (CNVs) are a type of polymorphism found to underlie phenotypic variation, both in humans and livestock. Most surveys of CNV in livestock have been conducted in the cattle genome, and often utilise only a single approach for the detection of copy number differences. Here we performed a study of CNV in sheep, using multiple methods to identify and characterise copy number changes. Comprehensive information from small pedigrees (trios) was collected using multiple platforms (array CGH, SNP chip and whole genome sequence data), with these data then analysed via multiple approaches to identify and verify CNVs.

**Results.:** In total, 3,488 autosomal CNV regions (CNVRs) were identified in this study, which substantially builds on an initial survey of the sheep genome that identified 135 CNVRs. The average length of the identified CNVRs was 19kb (range of 1kb to 3.6Mb), with shorter CNVRs being more frequent than longer CNVRs. The total length of all CNVRs was 67.6Mbps, which equates to 2.7% of the sheep autosomes. For individuals this value ranged from 0.24 to 0.55%, and the majority of CNVRs were identified in single animals. Rather than being uniformly distributed throughout the genome, CNVRs tended to be clustered. Application of three independent approaches for CNVR detection facilitated a comparison of validation rates. CNVs identified on the Roche-NimbleGen 2.1M CGH array generally had low validation rates with lower density arrays, while whole genome sequence data had the highest validation rate (>60%).

**Conclusions.:** This study represents the first comprehensive survey of the distribution, prevalence and characteristics of CNVR in sheep. Multiple approaches were used to detect CNV regions and it appears that the best method for verifying CNVR on a large scale involves using a combination of detection methodologies. The characteristics of the 3,488 autosomal CNV regions identified in this study are comparable to other CNV regions reported in the literature and provide a valuable and sizeable addition to the small subset of published sheep CNVs.

## Background

Copy number variants (CNVs) are a type of genomic polymorphism that potentially underlie a significant fraction of phenotypic variation [1]. CNVs are structural variants, defined as stretches of DNA that are greater than 1 kilobase (kb) in size and are duplicated or deleted in the genome of some individuals [2]. Mutation rate estimates for CNVs vary from 1.1×10^−2^ [3] to 1×10^−8^ per locus per generation [4, 5], which reflects the diverse processes by which CNVs are created. They can be over 1 megabase (Mb) [6] and are thought to comprise approximately 1% of an individual’s genome, which is much higher than the 0.1% thought to comprise SNPs [7, 8]. CNVs can be present in the same or overlapping regions of the genome in multiple individuals, these regions are called copy number variant regions (CNVRs). Copy number variants are distinct from another type of variant, indels (INsertions/DELetionS), in that indels are typically less than 1kb [2]. By definition they are also distinct from segmentai duplications (SD). Segmental duplications are defined as being over 1kb in length with at least 90% sequence identity between the duplicated segmants and are often not polymorphic in the population [9]. In many cases it is likely that Segmental duplications were once CNVs that have subsequently become fixed in the population.

There are many examples, particularly in humans, of CNVs influencing traits. These include multiple examples of CNVs associated with cancer susceptibility [10–12], the association of the FCGR3B gene copy number variant with systemic lupus erythematosus (SLE) [13], and CCL3L1 gene copy number, which has been linked to HIV susceptibility [14]. There is also evidence for CNVs influencing traits in other animal and livestock species. A 133kb duplication containing four genes causes hair ridge in Rhodesian and Thai Ridgeback dogs [15]. The chicken Peacomb phenotype is under sexual selection and is caused by a 3.2 kb duplication in an intron of the SOX5 gene [16]. The Peacomb allele contains ∼30 copies of the duplication, with variation in copy number present within individuals with the Peacomb phenotype. In pigs, Chen *et al* [17] found seven copy number variable genes that overlapped quantitative trait loci (QTL) for, among other traits, carcass length, backfat thickness, abdominal fat weight, length of scapular, intramuscular fat content of *longissimus* muscle, body weight at 240 days and glycolytic potential of longissimus muscle. Although not an association analysis, Chen *et al* [17] identified one CNV that had previously been associated with skin colour in pigs [18].

There have been many CNV studies in cattle, with a range of platforms used to identify CNVs [19–26]. Between 51 and 1265 CNVRs [20, 22] have been identified in the various cattle studies, with estimates of the proportion of the cattle genome thought to contain CNVRs ranging from 0.5 to 20% [22, 24]. Although the latter is likely to be an overestimate, the wide range in estimates is likely due to a number of factors, including the technology used to detect CNVs, different CNV calling criteria used, and the number of animals examined

While there is one notable example of a CNV having a direct effect on a sheep trait – the agouti duplication influencing coat colour [27] - to date, little work has been published on copy number variants in the sheep genome. An initial survey assayed eleven sheep on a cattle Roche-NimbleGen 385K oligonucleotide CGH array (oligo aCGH) which included 385,000 probes that were designed based on the cattle genome build btau_4.0 [28]. That study identified 135 CNV regions (CNVR) that covered approximately 0.4% of the sheep genome and ∼0.01-0.13% of each individual’s genome, which is substantially less than the approximately 1% estimated by Pang et al [8] in humans. This suggests many more sheep CNVs remain to be identified.

A number of approaches have been used to detect the presence of CNV. The main platforms are comparative genomic hybridisation (CGH) arrays [29–33], SNP arrays [34–37] and depth of coverage metrics applied to whole genome sequence data (e.g., [38–42]). Further, there are a variety of algorithms that can be used to analyse available resultant data. Perhaps the most widely used platform is array CGH, as it represents a cost-effective method to detect CNVs on a genome-wide scale in multiple individuals [43].

Trios have been used in CNV studies to determine the de novo mutation rate and to identify CNVs that represent heritable genetic units [4, 22, 44, 5]. This involves identifying CNVs in a fathermother-progeny trio. CNVs present in progeny and at least one parent are thought of as heritable and CNVs present in progeny but not in either parent indicate either a de novo mutation or an error in CNV identification. Given that CNVs are difficult to detect regardless of the platform or methods used, the best approach appears to be the conservative use of multiple methods to generate a set of high confidence CNV calls.

Given the lack of a comprehensive study of sheep CNVs, the objective of this study was to conduct a survey of sheep CNVRs using a range of detection methods. A Roche-NimbleGen 2.1M CGH array was designed and 36 animals (which included sets of trios) were assayed. Independent detection approaches were used in an attempt to validate the results. Finally, the CNVRs detected in this study were compared to those reported in an earlier survey of the sheep genome [28] and those detected in seven separate cattle studies [19, 20, 25, 28, 21–23].

## Results

### Roche-NimbleGen 2.1M CGH array construction and application

A total of four methodologies were used to detect CNV, with the main approach being the development and application of a 2.1M probe CGH array for the sheep genome. In total, 2,012,210 probes were designed with an average spacing across the autosomes of approximately 1.2 Kb. The array was used to assay a total of 36 sheep genomes, consisting of 30 individuals drawn from the International Mapping Flock [45] and a further six from a Reference Panel of International Sheep Genomics Consortium (ISGC) sheep (Supplementary Table 1). The Roche-NimbleGen segMNT algorithm was used to call CNV segments in each animal compared to the reference animal. Many different algorithms and criteria can be used to identify CNVs in array CGH data. Criteria employed to filter CGH data include restricting calls based on probe number within the CNV segment and log_2_ ratio (the ratio of test to reference probe intensity values) (Bickhart *et al* (2012); Liu *et al*., 2010; Fontanesi *et al*., 2011); Conrad *et al*., 2010); (Kijas *et al*., 2011)). These criteria are often selected using the results from a self-self hybridisation experiment, whereby self-self calls are used to indicate false positive calls, and rely on the assumption that the self-self hybridisation CNV calls cover the range of characteristics of false positive calls. It requires selecting a balance between filter values for number of probes and log_2_ ratio, so as to eliminate self-self hybridisation calls and other false positives from the dataset. Other studies have used differences between expected *versus* observed probe intensities on the sex chromosomes to set log_2_ ratio filters (Conrad *et al*., 2010). However, this does not account for possible probe number differences between true versus false CNV calls. As well as these filtering criteria, trios can be used to identify CNVs (Abecasis *et al*., 2010; Kijas *et al*., 2011; Krumm *et al*., 2012; Michaelson *et al*., 2012).

In this study, rather than using self-self hybridisation results to empirically set filters to remove false positives, a combination of trios and self-self hybridisation results were used to develop a logistic regression model for predicting whether or not a CNV segment represented a true CNV. The logistic regression model was developed using known positives (trio calls) and known false positives (self-self hybridisation calls) and the following variables were tested to determine if they were significant in predicting true versus false CNV segment calls: absolute log_2_ ratio of the CNV segment call; whether the call was a deletion or duplication; length of the call (base pairs); natural log transformed length variable; double natural log transformed length variable; the square of the length variable; number of probes in the CNV segment call; natural log transformed probe variable; double natural log transformed probe variable; the square of the probe variable; and corresponding two‐ and threeway interactions. The variables that were significant in predicting true versus false CNV segment calls were the absolute log_2_ ratio of the CNV segment call, the double natural log transformed length variable and the double natural log transformed probe variable. The resultant model was then used to predict true CNVs in the wider dataset, with some further downstream processing. The total number of autosomal segment calls predicted to represent true CNVs by our model, using CGH data from 30 animals, was 12,802. After removing calls based on a series of quality filters, a total of 9,789 autosomal CNV calls remained (Table 1). The mean absolute log_2_ ratio of these calls was 0.54 and the average length was 30kb with a range in length of 1kb-2.5Mb (Table 1).

**Table 1.** Characteristics of CNVs predicted true by the model (n=9,789) and filtered to remove artefacts.

| Variable | Mean | Median | Std dev | Min | Max |
| --- | --- | --- | --- | --- | --- |
| absl2r* | 0.54 | 0.43 | 0.36 | 0.25 | 3.47 |
| length (bp) | 30,332.02 | 8,706 | 107,369.37 | 1,003 | 2,522,449 |
| Datapoints<sup>#</sup> | 14.99 | 9 | 23.91 | 3 | 446 |

\*absl2r is the absolute log<sub>2</sub> ratio of the CNV. # number of CGH array probes in the CNV.

On average, 326 CNVs were detected per individual, with a median of 321 and range of 109 to 643. One animal had notably more CNV calls than the other animals, however, it had the same CNV content on the autosomes (as a percentage of total length in base pairs) as the other animals.

### Autosomal CNVR

CNV information from all animals was combined to obtain 3,488 CNV regions on the ovine autosomes (Supplementary Table 2). The average length of these CNVRs was 19kb, with a range of 1kb to 3.6Mb. Shorter CNVRs were more frequent than longer CNVRs in the genome. The total length of all CNVRs was 67.6Mbps, which equates to 2.7% of the sheep autosomes. For individuals, this value ranged from 0.24 to 0.55%. Most CNVRs were seen in just one animal (Figure 1), however 1,424 (41%) were independently called in at least 2 individuals. A small percentage (0.11%) of CNVRs were observed in all animals, which likely indicates the presence of a CNV in the reference animal only - the ‘reference effect’ [46]. The majority of CNVRs (58%) contained only deletion CNVs, 38% of CNVRs contained only duplication CNVs and 4% were compound CNVRs, containing both duplication and deletion CNVs.

**Table 2.**
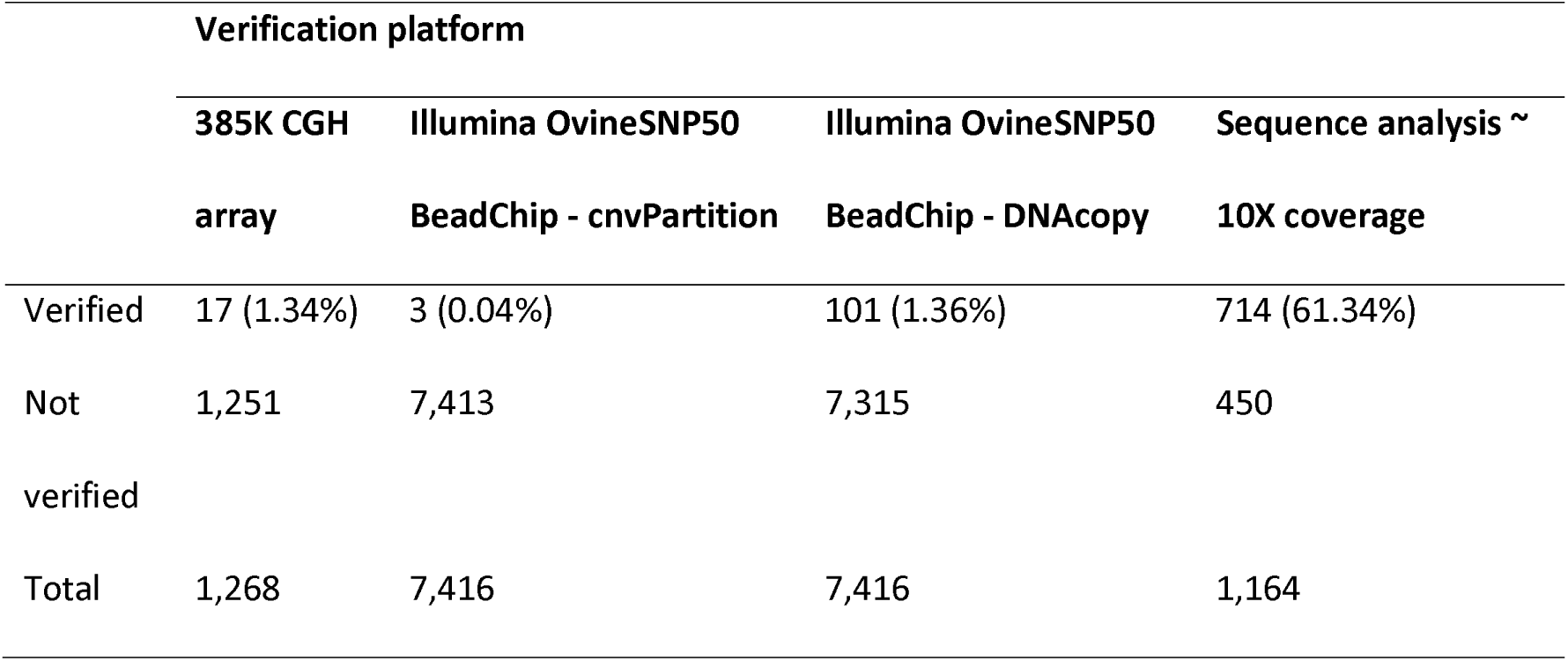
Characteristics of CNVs predicted true by the model (n=9,789) and filtered to remove artefacts.

**Figure 1.**
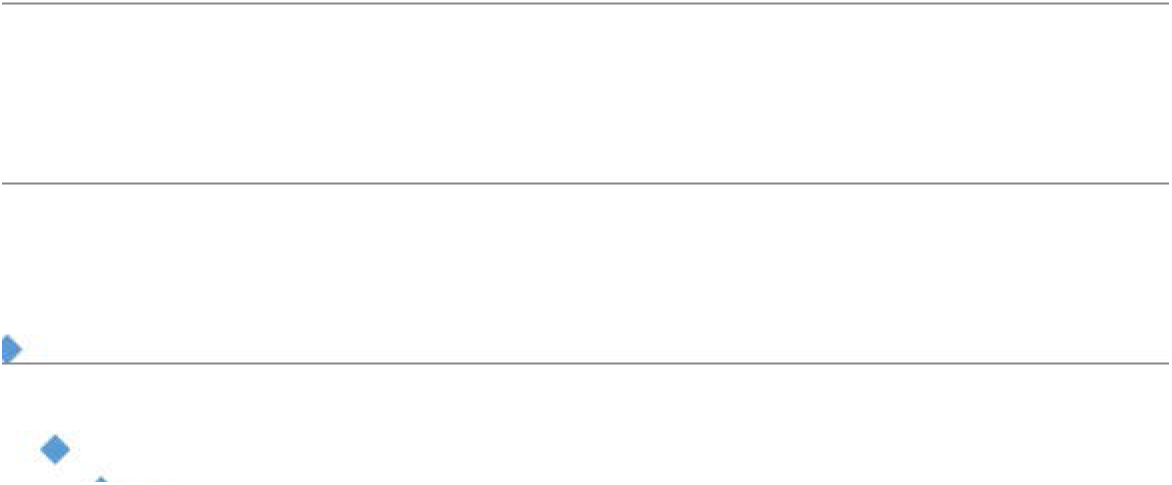
CNVR frequency across animals.

The number of CNVRs on each chromosome ranged from 76 on chromosome 27 to 185 on chromosome 19 (Figure 2). As can be seen in Figure 2, there was a weak positive linear relationship between chromosome length and number of CNVRs (R^2^=0.27).

**Figure 2.**
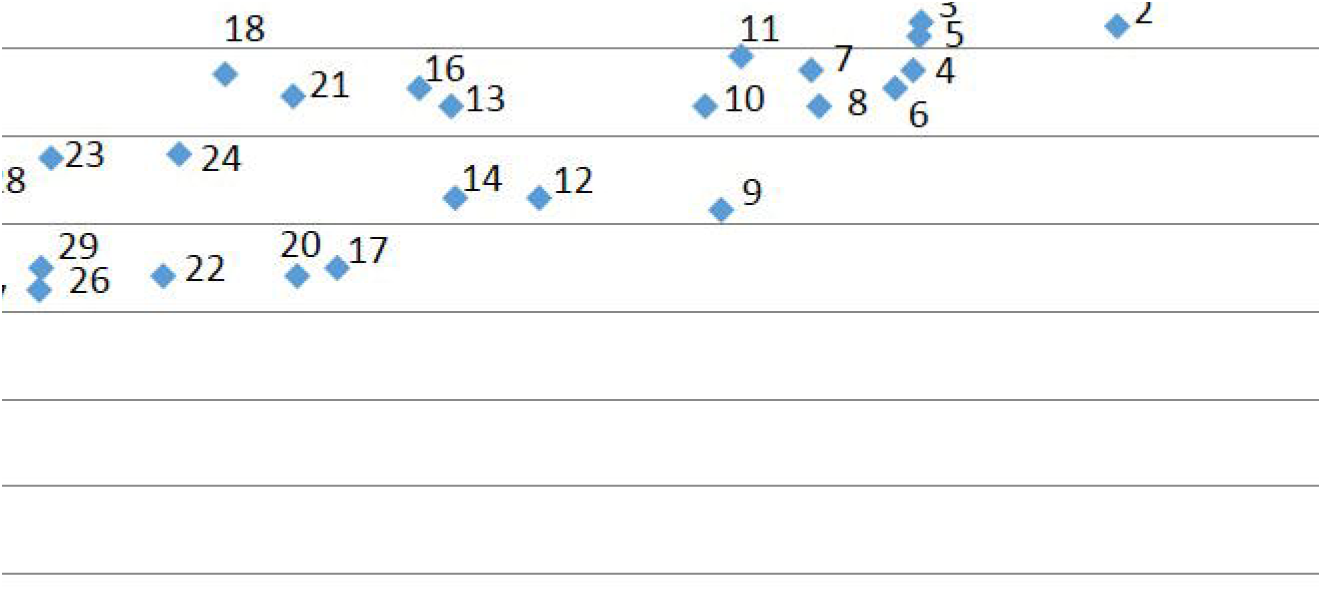
Number of CNVRs by chromosome length. Labels correspond to chromosome number.

The average spacing between CNVRs ranged from one every 347kbp on chromosome 19 to one every 1.2Mb on chromosome 1. The closest CNVRs were approximately 1.5kb apart, while the largest distance separating CNVRs was 8.5Mbps. The two-sample Kolmogorov-Smirnov test showed that the distribution of the CNVRs in the genome (in terms of the inter-CN V distance) was significantly different to that which would be expected should the CNVRs be uniformly distributed (p-value = 4.56xl0^−7^). Specifically, the CNVRs tended to be clustered together in the genome (Figure 3).

**Figure 3.**
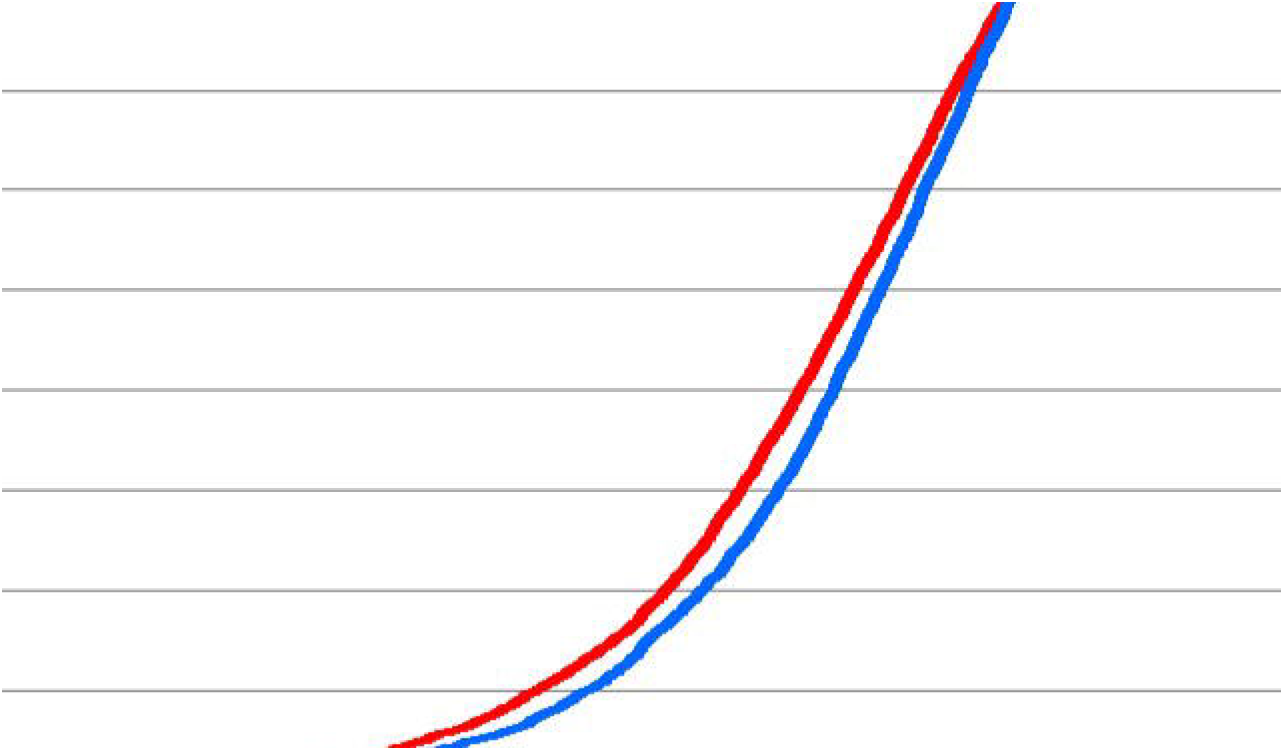
Cumulative density plot of the distances separating CNVRs. The red line reflects the observed pairwise distances between CNVRs, while the blue line reflects the simulated (expected if CNVRs are uniformly distributed in the genome) distances separating CNVRs.

### Cross platform verification of autosomal CNVRs in sheep

A small subset of animals assayed with the 2.1M CGH array were also used for data generation with a lower density 385K CGH array (5 individuals), the OvineSNP50 BeadChip (24 animals) and whole genome sequence from the six reference panel animals (Supplementary Table 1). This facilitated an examination of the proportion of CNVRs independently called across platforms.

The verification rate of CNVRs identified on the 2.1M CGH array on both the 385K CGH array and the OvineSNP50 BeadChip was low. Results from these analyses are presented in an additional file [see Additional file 1].

The final comparison utilised analysis of whole genome sequence from the six reference panel animals. Each individual was sequenced to between 9.8X and 14X genome wide coverage before variation in read depth was used to detect CNVR (see Methods). The same six animals had 852 CNVRs arising from 1,164 CNV calls detected using the 2.1M CGH array. Comparing the CNV calls revealed 61% of the Roche-NimbleGen 2.1M CGH array CNV calls were independently identified in the sequence data (Table 2). Two thirds of the CNV calls that were verified were observed as a consistent deletion or duplication CNV across platforms in a specific animal. The remaining verified CNVs were observed as a CNV of the opposite type (deletion versus duplication) in the Poll Dorset animal. This animal was used as the reference animal on the Roche-NimbleGen 2.1M CGH array and therefore CNVs in this animal can be incorrectly observed as CNVs in the test animal when in fact no CNV is present in the test animal. That is, a deletion in the Poll Dorset may be observed as a duplication in the test animal on the 2.1M CGH array, while in the sequence data, the test animal shows no CNV in the region but the Poll Dorset shows a deletion. The same is true for duplications in the Poll Dorset, which will be observed as deletions in the test animal, even if no CNV is present in the test animal in that region.

There were instances where the sequence data showed that there was a CNV in the Poll Dorset and the test animal in the same region, but the type (duplication/deletion) of CNV in the test animal was not consistent between the 2.1M CGH array and sequence platforms. For example, a 2.1M CGH deletion that was observed as a duplication in the test and reference animal in the sequence data. These calls were considered to be verified as there were still CNVs present in the sequence data and it is possible that the magnitude of the log_2_ ratio of the CNV call on the 2.1M CGH array was higher in the Poll Dorset than the test animal which could result in inconsistencies between the types of CNVs detected. There were instances in the data where a CNV call of one particular CNV region could be verified in one animal and not in another animal, which indicates that the CNV is likely present in both animals but the sequence analysis failed to identify the CNV in one of the animals.

Significant differences in absolute log_2_ ratio, length and GC content were observed between the sequence verified and non-verified 2.1M CGH array calls. Verified calls had higher absolute log_2_ ratios (0.62 versus 0.50) and were longer (46kb versus 9kb) on average than non-verified calls. This suggests that longer calls with higher absolute log_2_ ratios are either more likely to represent true CNVs or are easier to verify than shorter calls with lower absolute log_2_ ratios. Sequence corresponding to non-verified calls showed significantly higher (two-tailed t-test for proportions) GC content on average compared to verified calls –44.6 versus 43.0%. Both verified and non-verified calls had significantly higher GC content compared to the genome average (42.6%). More duplications (72.4%) than deletions were verified on the sequence platform - 72.4% versus 54.7%. This is not surprising, as there was less variation in the sequence data in regions with low read depth, which reduces the ability to detect differences in copy number in these regions and hence also CNVs relating to deletions.

### Comparison of autosomal CNVRs to those identified in the sheep and cattle literature

In total, we detected 378 (18%) of the 2,154 CNVRs reported in seven other sheep and cattle studies. Of the 2,154 CNVs detected in the seven other studies, 352 were present in more than one study.

We detected 132 (38%) of the 352 CNVs observed in multiple studies, whereas we only detected 14% of the CNVRs observed in just one other study (Table 3). The more frequently a CNVR was observed in the other studies, the more likely we were to detect the CNVR (Table 3). We were able to detect 31% of the CNVRs identified in the initial sheep study by Fontanesi *et al* [28] and between 16-62% of CNVRs detected in the cattle studies.

**Table 3.** Comparison between CNVRs observed in this study and CNVRs observed in the literature.

| Number of studies CNVR<br>observed in | Number of CNVR | Number of these CNVR identified in this<br>study (%) |
| --- | --- | --- |
| 1 | 1,802 | 246 (13.7) |
| 2 | 255 | 82 (32.2) |
| 3 | 66 | 24 (36.4) |
| 4 | 20 | 16 (80.0) |
| 5 | 7 | 6 (85.7) |
| 6 | 4 | 4 (100) |

Eleven percent of the 3,336 CNVRs detected in this study and successfully mapped to the btau_4.0 genome overlapped CNVRs in these other studies. This is lower than would be expected based on overlap between CNVRs from the other studies with each other, which ranges from 20-77%. By comparison, 28% of the CNVRs from the sheep study by Fontanesi *et al* [28] were observed in at least one of the cattle studies.

### Overlap between autosomal CNVRs and genes

Of the 3,335 CNVRs identified on the Roche-NimbleGen 2.1M CGH array that mapped to OARv3 autosomes, 1,335 (40%) overlapped the coding sequence of one or more genes; 45% of duplication CNVRs, 36% of deletion CNVRs and 59% of deletion/duplication CNVRs overlapped genes. The proportion of duplications overlapping the coding sequence of genes was significantly different (Chisquared test, p < 0.0001) to the proportion of deletions overlapping genes. Based on permutation analysis, these proportions were significantly greater than that which would be expected if the CNVRs were randomly distributed in the genome (p=0.01). Both the agouti signalling protein and adenosylhomocysteinase genes were overlapped by one of our CNVRs, which confirms the presence of the agouti duplication reported by Norris and Whan [27] in this dataset, and thus provides a positive control for the CNVR identification methods presented here. It is important to note that the agouti duplication can be present in multiple copies [27], hence the reason that it shows up even upon comparison to another white fleeced sheep.

### Non-autosomal CNVRs

The total number of chromosome X Roche-NimbleGen 2.1M CGH array segment calls predicted to be real was 697, however, 308 of these were observed as deletions in males. It is possible some of these are real, particularly if they are present in the pseudo-autosomal region, however, this cannot be confirmed in our analysis as we do not have a clear pseudo-autosomal boundary defined. After filtering all 697 CNV calls based on size and log_2_ ratios, 615 of these were predicted to be real, however, only 317 were either deletions or duplications in females or duplications in males. These 317 were used to call CNVRs on chromosome X. In total, we estimate there are at least 114 CNVRs on chromosome X, representing approximately 3.2% of the length of the X chromosome. In addition to chromosome X CNVRs, four CNVRs were identified on UMD3_OA_chrun, observed in one to ten animals. These CNVRs spanned a total length of 19,304bps.

Including the 3,488 CNVRs observed on the autosomes, the 114 CNVRs observed on chromosome X and the 4 CNVRs identified on chromosome unknown (UMD3_OA_chrun), we estimate there to be approximately 3,606 CNVRs in the sheep genome. This includes CNVRs identified on chromosome X and UMD3_OA_chrun. The total length of these 3,606 CNVRs is estimated to be 72.4Mbps, however, it is possible that some of the CNVRs on UMD3_OA_chrun may overlap those identified on the autosomes and therefore this number may be slightly lower.

## Discussion

The results reported here provide a genome wide view of the frequency of CNV, an important class of genomic variant that is currently poorly characterised in the sheep genome. Using a custom built Roche-NimbleGen 2.1M CGH array, 9,789 autosomal CNVs were detected in 30 sheep. On average these CNVs covered 0.4% of each animal’s genome. This is higher than that reported in the initial sheep survey where, on average, 0.05% of an individual sheep genome comprised CNVs [28]. The difference in estimates is not surprising as this study used a CGH array with 2.1 million probes while Fontanesi *et al* [28] used a CGH array with 385,000 probes. Based on probe spacing in the genome and the filters applied to the data, the earlier study detected CNVs greater than 30kb in length, on average, while this study had a resolution of ∼4kb on average. As a result, differences in resolution may have resulted in differences in the number of CNVs detected. This is reflected in the datasets, with the average size of CNVs detected by Fontanesi *et al* [28] being 77.6kb (median 55.9kb) and the average size detected in this study being 30.3kb (median 8.7kb). The individual genome CNV composition estimates are similar to, but slightly lower than, estimates reported in humans (e.g., 0.5%, [48]; 0.78%, [7]; and 1.2%, [8]).

The 9,789 autosomal CNVs reported in this study correspond to 3,488 autosomal CNV regions in the 30 animals tested, representing 2.7% of the sheep genome. This is approximately seven times higher than estimated in the initial sheep survey [28], which is to be expected as more animals were assayed in this study. This estimate is similar to the range of estimates in cattle [19, 25, 21, 26, 22, 23, 20] and again similar but slightly lower than estimates in humans (3.7%, [7]; 5%,[48]). Estimates in humans are likely to provide a more accurate estimate of CNV composition in the genome, as studies have involved more individuals and used a wider range of technologies, often employed together. As in the Fontanesi *et al* [28] study, this study suffers from the lack of a complete reference sheep genome. We used a sheep genome that was constructed using a cattle reference genome to design probes for inclusion on the 2.1M CGH array. The genome used, UMD3_OA, does not include any regions that are present in the sheep genome but that are not present in the cattle genome. This means that sheep CNVs in regions deleted or of low homology in the cattle genome are likely to have been undetected in this study. Future work will benefit from using a sheep reference genome for CNV analysis. However, the CNVRs presented in this study provide a substantial addition to the currently published sheep CNV regions, and bring the resource up to a level similar to that available in cattle.

There were also 118 CNVRs identified on chromosome X and chromosome unknown. However, these were lower confidence calls and were not considered in further analyses. Of the 3,488 autosomal CNVRs identified in this study, 59% were observed in just one animal, which is comparable to results in the literature [7, 35, 22, 23, 37]. One and a half times more deletions than duplications were observed. This imbalance is one that is commonly reported in the literature [49, 50, 22] and could be due to ascertainment bias. The ascertainment bias arises because the proportional difference between probe intensity of test and reference animals is greater for copy number losses than gains meaning that deletions are easier to detect than duplications.

The CNVRs detected in this study tended to be clustered together in the genome. This may be an artefact of the segMNT algorithm and our CNVR calling algorithm, which may have failed to collapse multiple CNVRs originating from one CNVR into one region. However, similar distributions have been reported in other studies [5, 51–53] and also for the closely related Segmental duplication variant [9]. If this clustering represents the true underlying distribution in the genome, then it may indicate that the clustered CNVRs are the result of increased mutational activity in repetitive regions of the genome which could facilitate mechanisms such as non-allelic homologous recombination [54]. Determining if the CNVRs are a result of one mutational event or multiple mutational events would require detailed analysis of specific regions, probably using deep sequencing.

There are reports in the literature that CNVRs are preferentially located outside of gene regions [51, 55, 56, 37] and that those CNVs that do overlap genes are more likely to be duplications than deletions [7, 57, 37]. The rationale is that deletions are more disruptive to gene function than duplications and therefore are subject to greater selective pressure. In this study, a significant difference was observed in the proportion of duplications overlapping the coding sequence of genes compared to deletions –0.45 versus 0.36. However, both of these proportions were significantly higher than would be expected if CNVRs were randomly distributed throughout the genome. Therefore, in this study there is no evidence to suggest that the CNVRs identified in this study are preferentially excluded from genic regions as has been suggested in the literature. Other results reported in the literature have also found an enrichment of CNVs in these regions [30, 53]. Cooper *et al* [53] suggest that CNVs that overlap Segmental duplications (SDs) are more likely to be enriched in genic regions, while CNVs that do not overlap SDs are enriched in gene poor regions of the genome. As genes and Segmental duplications are GC rich [58] and GC rich regions are more prone to CNV formation, then it is possible that certain types of CNVs are enriched in genic regions. While selection against or for CNVs and CNV formation mechanisms are reasonable explanations for the depletion or enrichment of CNVs in genic regions, it is also possible that differences reported in the literature are due to ascertainment bias introduced by using different methods for CNV detection. Again, this illustrates the difficulties associated with CNV identification.

Compared to the lower density 385K CGH array and the OvineSNP50 BeadChip, whole genome sequencing exhibited the highest cross platform verification rate, with 61% of CNVs verified with this platform. The CNVs that were unable to be verified were shorter and had lower absolute log_2_ ratios than calls that were able to be verified. Both verified and non-verified CNVs had significantly higher GC content than the genome average, which supports data from the literature reporting that GC-rich regions can be more prone to CNV formation [61, 62]. Non-verified CNVs had significantly higher GC content than verified CNVs. While it is possible that the non-verified CNVs were false negatives in the sequence analysis, it is also possible that they were false positives in the CGH dataset, as false positive CGH calls can be related to regions with high GC content [63, 64]. Future work could involve adjusting CGH intensity data for GC content.

This study detected 18% of the CNVRs reported in seven other sheep and cattle studies [19, 20, 25, 28, 21–23]. Thirty one percent of the CNVRs that were previously detected in an initial survey of CNVs in the sheep genome [28] were detected in this study. We were able to identify all of the CNVRs that were observed in six of the other studies, but only 14% of CNVRs observed in just one other study. In fact, the more studies a CNVR was detected in, the more likely we were able to identify the CNVR in our analysis. This trend was also reported by Kijas *et al* [22]. This suggests that either these CNVRs are less likely to be false positives or they may be more common than the CNVRs detected in just one study or, alternatively, they may be more likely to occur in both sheep and cattle. Common CNVRs will be present in more individuals in the population and therefore are more likely to be observed in the diverse range of animals tested in the different studies. Reasons that this study was unable to detect many of the CNVs from the other studies include: CNVs that occur in cattle but not sheep; rare CNVs not seen in our sample of sheep; and false negatives in our study due in part to the different methods used for CNV detection. Similarly, only a small number (11%) of CNVRs identified in this study overlapped CNVs detected in these seven other studies. Again, lack of overlap could be due to the different species or individual animals tested, different methods used for CNV detection, false negatives in other studies and false positives in our dataset. Confirmation rates varied widely across the studies compared to our results. Variation in confirmation rates from different studies has also been reported in the literature for human CNV studies [66, 67].

## Conclusions

In this study, comprehensive information from trios, multiple platforms and different algorithms were used with the aim of verifying CNV segment calls from the Roche-NimbleGen 2.1M CGH array. CNVs are difficult to verify and as is observed in the literature, a combination of approaches appears to be the best way to accurately detect CNVs on a large scale. It is likely that comprehensive sequencing or qPCR would provide clearer information about individual CNV regions and give an indication of the accuracy of the methods used to detect them. Regardless, characteristics of the CNV regions detected in this study are comparable to those reported in the literature, and the CNV regions identified here add to the initial survey of CNVs in the sheep genome by Fontanesi *et al* [28].

## Methods

### Roche-NimbleGen 2.1M CGH array - design overview

In total, 2,012,210 probes (50-75 base pairs in length) were distributed evenly on non-repetitive regions of the UMD3_OA ovine genome build (an in-house AgResearch comparative sheep genome assembly, built using cattle reference genome UMD3 [68] and accessible at www.sheephapmap.org/CNV/), with an average spacing of approximately one probe per 1,250 base pairs (bps) on the autosomes and one probe per 1700bps on chromosome X. In addition to these probes, a further set of probes was designed around SNPs found on the Illumina OvineSNP50 BeadChip, with the aim of increasing cross platform validation between the 2.1M CGH array and OvineSNP50 BeadChip. This involved mapping SNPs and flanking sequence onto UMD3_OA. In some instances, SNP sequences did not map uniquely to the genome, with multiple hits on the same chromosome, suggesting the possibility that multiple copies of the sequence could occur in adjacent duplicated regions (e.g. CNV). As these SNPs may have been in CNV regions, these regions were also used for specific probe design and inclusion on the array. Probes were also designed on chromosome unknown scaffolds. Chromosome unknown scaffolds represent sequence data that cannot be placed on the genome assembly.

### Roche-NimbleGen 2.1M CGH array design - targeted probe design around OvineSNPSO BeadChip SNPs

In total, 28,754 out of 50,064 SNP sequences (either the 50bp OvineSNP50 BeadChip probe or 300bp flanking the SNP) successfully mapped to UMD3_OA (BLAST parameters ‐U T-F "m D" ‐e 1e-5, Korf *et al* [69]) and met the requirement of having three probes designed to cover them, as selected by one of the following two methods (Figure 4). The first involved designing a probe to cover the SNP base pair position. Flanking probes were designed within 400bp windows 100bp up‐ or down-stream of the SNP region, where the SNP region consisted of 300bps flanking the SNP position. If three probes were not obtained with this method, then a second method was used. This involved selecting a probe in the SNP region without requiring the probe to cover the SNP position, with flanking probes selected from 400bp windows 100bp up-or down-stream of the SNP region (Figure 4). In total, 86,262 probes were designed within or adjacent to 28,754 SNP regions.

**Figure 4.**
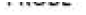
Selection of CGH array probes to cover OvineSNP50 BeadChip SNP positions. Two methods were used to select probe sets to cover SNPs. The first method (a) involved designing at least one probe to cover the SNP position, with two probes in flanking regions. The second method (b) involved designing a probe within the 300bp region surrounding the SNP and two probes in flanking regions.

Of the 21,310 SNP sequences that could not be mapped to UMD3_OA, 240 were mapped by relaxing the BLAST parameters to ‐W 11 ‐q ‐1 ‐r 1 ‐s 0 ‐F "m D" ‐U T ‐X 40 [69]. A total of 634 probes were designed to cover 218 of these SNP regions.

A subset of 401 SNP sequences mapped to UMD3_OA, but not uniquely - with two top hits on the same chromosome. In total, 879 probes covering 323 of these positions were designed for inclusion on the 2.1M CGH array.

### Roche-NimbleGen 2.1M CGH array design - chromosome unknown

Chromosome unknown sequences (n=492) were merged into a Virtual chromosome, UMD3_chrll_OA, with each sequence separated by 100 N's. Probes were distributed at an average spacing of approximately one every 1,600bps on this chromosome.

### Roche-NimbleGen 2.1M CGH array – animals assayed

Genomic DNA was extracted from blood samples of 36 animals (Supplementary Table 1), which were assayed on the 2.1M CGH array. Thirty animals were from the International Mapping Flock (IMF) and consisted of families of trios (Figure 5). The IMF animals are crossbreds of up to five different breeds–Texel, Coopworth, Perendale, Romney and Merino [45]. In addition to the IMF animals, six sheep, sequenced to approximately 10X coverage each, were also assayed on the 2.1M CGH array. These six animals were - Awassi, Merino, Poll Dorset, Romney, Scottish Blackface and Texel purebreds. The Poll Dorset was used as the reference animal for all 2.1M CGH array hybridisations and was also run against itself in a self-self hybridisation to allow characterisation of false positive calls [70, 23].

**Figure 5.**
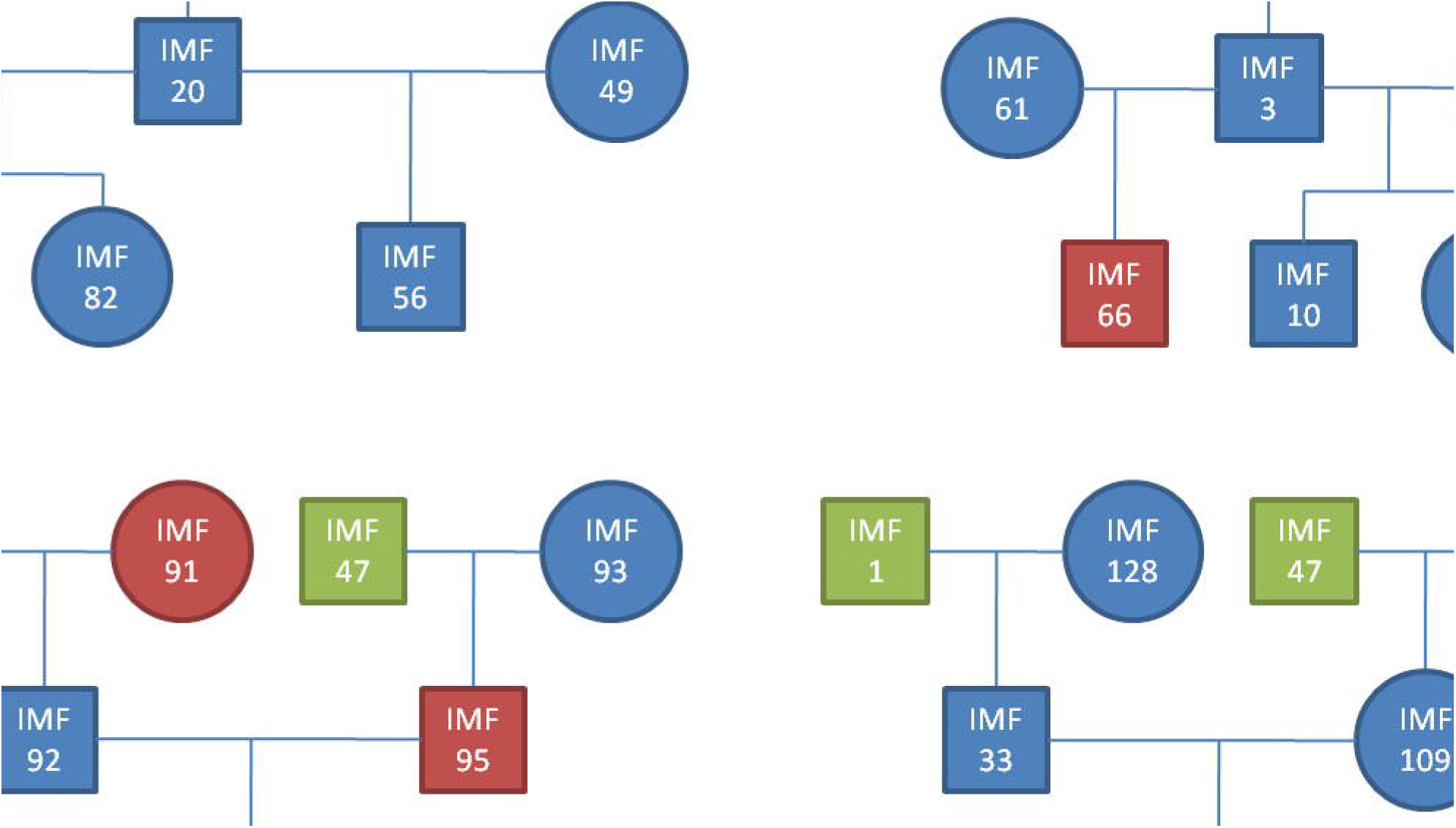
Pedigree of International Mapping Flock (IMF) animals assayed on the Roche NimbleGen 2.1M CGH array. Some animals (green) appear in more than one pedigree. Segment calls from animals IMF66, IMF91, IMF95, IMF108 and IMF112 (red) were removed from the analysis due to failed 2.1M CGH arrays.

### Roche-NimbleGen 2.1M CGH array - segMNT output processing

CNV segments were called in the assayed animals by Roche-NimbleGen using their proprietary segMNT algorithm. This software reports the average log_2_ ratio of a segment (the binary logarithm of the average of the intensity of the test animals probes in a segment call divided by the average of the intensity of the reference animals probes in the same region), the number of datapoints (probes) included in the segment and the length of the segment in base pairs.

The variance of normalised log_2_ ratio values over all probes for each animal was obtained. Five animals were deleted from the analysis as their log_2_ ratio data exhibited larger variation than observed in other animals, meaning that they were deemed to be failed CGH hybridisations.

Segment calls with absolute log_2_ ratios less than 0.1 were removed from the analysis [7].

### Validating Roche-NimbleGen 2.1M CGH array segment calls

IMF trios were used to validate segment calls. If a progeny segment call was seen in at least one parent at an identical genomic location (same first and last probe included in the segment call and therefore same genomic start and stop position), the progeny call was considered validated. These calls were deemed to represent “true CNVs” for model building.

### Model used to predict CNVs in the wider dataset and downstream filtering

For model building, validated progeny calls were deemed to represent true CNVs and self-self hybridisations were deemed to be false positives. Only autosomal segment calls were used. Forward stepwise logistic regression was used to construct a model, with a binary outcome variable 0 (self-self) or 1 (validated trio segment call). Variables used for model building were: absolute log_2_ ratio (absl2r); whether the call was a deletion or duplication; length, in bps; ln(length); ln(ln(length); length-squared; number of probes in segment call, datapoints; ln(datapoints); ln(ln(datapoints); datapoints-squared; and corresponding two-and three-way interactions. If the Wald chi-square statistic for a variable was significant at the 0.3 level it was added to the model. A variable remained in the model if it was significant at the 0.35 level.

The crossvalidate procedure in SAS software (*SAS version 9.1*) was used to test model performance. This procedure omits one segment call in turn and re-calculates model coefficients based on all other segment calls per iteration. It then predicts the probability the omitted call represents a true CNV. Threshold values were applied to categorise calls as true or false based on their probabilities - true or false. Probability thresholds tested were 0.5, 0.6, 0.7, 0.8, 0.9, 0.95, 0.96, 0.97, 0.98 and 0.99. For each probability threshold tested, the number of times the procedure correctly predicted the known segment call status (true or false) was used as a measure of model accuracy. The final probability threshold used was 0.95.

The final model selected was,

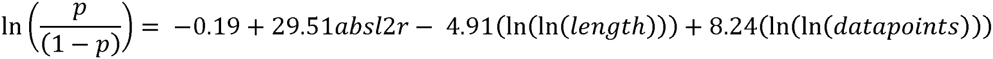

This model was applied to all segment calls not used in model development. Segment calls equal to or greater than the probability threshold of 0.95 were retained. The dataset was further filtered to include only CNVs >=1kbp in length (so that they conformed to the definition of a CNV, as per [2], only CNVs with >= 3 probes in the corresponding segment call and with absolute log_2_ ratio >=0.25. These filtered segment calls were deemed to represent true CNVs.

Segment calls on chromosome X were processed through the model and filtered as above. Filtered CNVs on chromosome X were considered to represent true CNVs for female individuals. Duplications on chromosome X in males were considered to represent true CNVs. Deletions on chromosome X in males were assumed to be inconclusive as they could be due to differences in the number of X chromosomes between the male test animal and the female reference animal.

Segment calls on the virtual chromosome UMD3_chrU_OA were processed differently to segment calls on the autosomes and chromosome X. Chromosome unknown sequences were collated into larger virtual chromosomes, UMD3_chrU_OA, with each sequence separated by 100 N's. Segment calls on this virtual chromosome were discarded if they spanned more than one chromosome unknown sequence or if all probes on one chromosome unknown sequence were included in the segment call. The reason for excluding segment calls where all probes on the chromosome unknown sequence were included in the call was because there was no way to compare the call to nearby sequence to determine if the log_2_ ratio was different to other stretches of DNA in the region. There were two Poll Dorset (self-self hybridisation) segment calls on UMD3_chrU_OA. The log_2_ ratios of these calls were ‐0.32 and ‐0.17. Thus calls with absolute log_2_ ratios ≤0.32 were removed from the analysis. Segment calls that met these criteria and that contained at least two probes, while excluding at least two probes from the corresponding chromosome unknown sequence, were retained.

### CNV regions

Across all animals, autosomal and chromosome X CNVs within 1,500bps of one another were collapsed into CNV regions (CNVRs).

To determine if CNVRs were uniformly distributed in the genome, a simulated dataset of CNVRs was generated by randomly sampling genomic positions of the identified autosomal CNVRs from a uniform distribution. Spacing was constrained so that CNVRs could not be within 1,500bps of each other. The simulated dataset provided an expected distribution of CNVRs in the genome and corresponding pairwise distances between CNVRs. A Kolmogorov-Smirnov test was performed to determine if the distribution of pairwise distances between CNVRs in the observed dataset was significantly different from that seen in the simulated dataset.

### Verifying CNVRs across platforms

Three other platforms were used for CNV identification – Roche-NimbleGen 385K CGH array, OvineSNP50 BeadChip, and Illumina HiSeq 2000 sequence data analysis, with each based on a different version of the ovine genome. To perform cross platform validation autosomal CNVRs identified on the Roche-NimbleGen 2.1M CGH array were mapped to genomes BTA_OARv.2 (for use with the 385K CGH array), OARv1 (for use with the OvineSNP50 BeadChip) and OARv3 (for use with sequence data analysis). CNVR sequence and 1,750bps flanking the start and stop of each CNVR were obtained. Sequences were masked with an ovine repeat database isgcandrepbase2 (Supplementary file 1) and BLASTed against each genome, with parameters ‐F 'm D' ‐U T ‐Z 2000 [69]. CNVR start and stop positions on each genome were approximated based on the BLAST alignment. When the predicted CNVR start position was a negative number, it was set to one (i.e. the first base pair of the chromosome).

The Roche-NimbleGen 385K CGH array is based on the same technology as the Roche-NimbleGen 2.1M CGH array; however, it has fewer probes covering the genome, with a probe density of approximately 1 probe per 6,000bps. Twenty animals were run on the 385K CGH array, including five animals (Awassi, Merino, Romney, Scottish blackface and Texel) that were run on the 2.1M CGH array. The Poll Dorset was used as a reference on the 385K CGH array and the 2.1M CGH array. Autosomal CNVRs identified using the 2.1M CGH array were positioned on BTA_OARv.2 as described above. CNVRs positioned on BTA_OARv.2 autosomes were retained for cross platform verification. CNV segments called by the NimbleGen segMNT software in the 385K CGH dataset were processed to include only autosomal segments with absolute log_2_ ratios ≥0.25. Autosomal CNVRs in the five animals were considered verified if there was overlap between their processed 385K CGH segment calls and their 2.1M CGH array CNVR calls mapped to BTA_OARv.2. This comparison was performed separately for each animal.

Twenty IMF and five sequenced animals had previously been genotyped on the OvineSNP50 BeadChip. SNP genotypes for these animals were run through the cnvPartition (Illumina Inc., USA) and DNAcopy [47] algorithms. DNAcopy results were filtered to include only calls with absolute log_2_ ratios ≥0.25. Autosomal CNVRs identified with the 2.1M CGH array and successfully mapped to OARv1 autosomes were considered verified if they overlapped autosomal CNVs predicted by cnvPartition or DNAcopy, in the same animal.

Six animals assayed on the 2.1M CGH array were each sequenced to between 9.8X and 14X coverage by paired-end sequencing on the Illumina HiSeq 2000 platform at Baylor College of Medicine (NCBI short read archive accessions SRX150284, SRX150292, SRX150299, SRX150330, SRX150341, SRX150350). The following analysis was carried out separately for each animal. Sequence reads were positioned on ovine genome OARv3 using the Burrows-Wheeler Alignment (BWA) algorithm [71] and pileup files [72] were used to retrieve read depth information at each base pair position on the autosomes. Reads were portioned into 1kbp overlapping bins, excluding repetitive sequence, using a sliding window of 200bps. Masked repetitive sequence positions were translated to genome build OARv3. As well as excluding repetitive sequence, for each chromosome a maximum read depth was set per chromosome to exclude potentially unmasked repeats from the CNV sequence analysis. The maximum read depth threshold was set based on inspection of the read depth distribution function with the aim of excluding outliers in read depth data. Bins with a maximum read depth exceeding the threshold were deleted from the analysis. The average read depth over all base pairs was determined for each bin after correcting for GC content based on methods presented by Yoon *et al* [73].

Pseudo-Maximum likelihood was used to fit a mixture model to determine if the average read depth for each bin represented a homozygous deletion (copy number, CN=0), heterozygous deletion (CN=1), normal diploid copy number (2), heterozygous duplication (3) or homozygous duplication (4) in the genome. The mixture model used (Table 4) was a mixture of four normal distributions (for modeling CN = 1 to 4) and one half-normal distribution (for CN = 0). Constraints were placed on the Parameters of the normal distributions so that the means and variances of the distributions corresponding to CN =1, 3 and 4 were equal to respectively 1/2, 3/2 and 2 times the mean and variance of the distribution corresponding to CN = 2. Model fitting was done on a per chromosome basis, using the R function *nlminb* [74]. Specifically, seven parameters were estimated for each chromosome: *μ*_2_ and 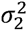 the mean and variance of read depth for a bin corresponding to CN = 2 (the “normal” diploid copy number); 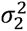 the variance of read depth for a bin corresponding to CN = 0 (homozygous deletion) and four of the five mixture weights (prior probability of a bin falling into each of the five distributions). Where these parameters could not be estimated for a chromosome, average estimates based on all other chromosomes for a given animal were used. Table 5 details the starting values and lower and upper bounds used by *nlminb* for each parameter. Based on those parameter estimates, each bin was assigned to one of the five CNV classes by multiplying the values of each of the five probability density functions for each bin by the corresponding mixture weights (i.e., calculating the posterior probability of a bin being in each of the distributions) and selecting the CNV class with the highest value. For each of the six animals, bins in regions corresponding to autosomal CNVRs identified on the 2.1M CGH array and mapped to OARv3 autosomes were used to determine if the CNVR was verified in the sequence data. Specifically, if at least one bin was observed as representing a CNV then the CNVR was considered to be verified. In instances where there was conflict between results from the sequence analysis and the 2.1M CGH array, individual animal data were compared to the reference (Poll Dorset) animal. This animal was used as the reference animal in the 2.1M CGH array experiments and therefore results for individual animals may be influenced by the corresponding copy number present in the Poll Dorset.

**Table 4.**
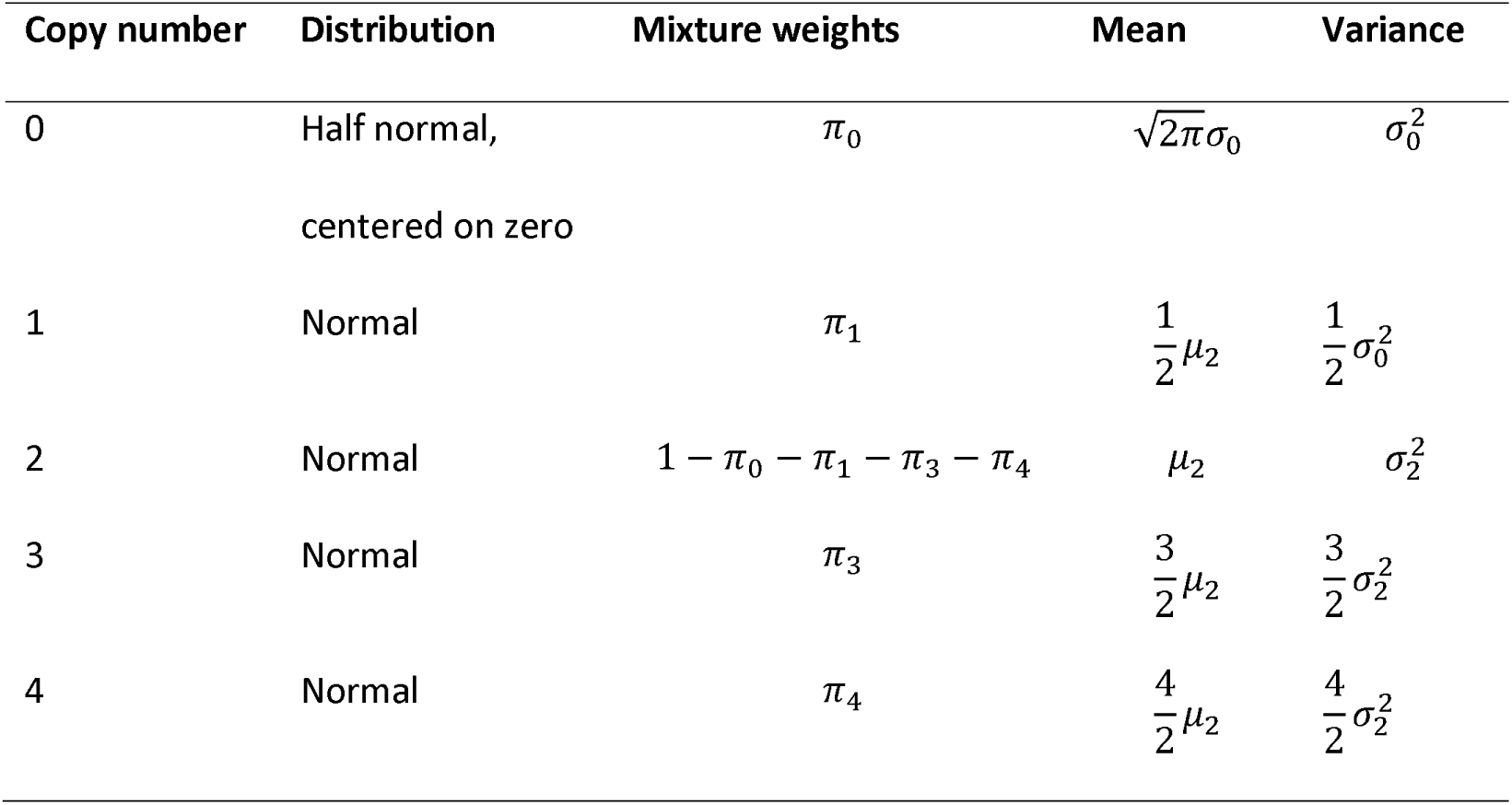
Description of the pseudo-maximum likelihood derived mixture model for estimating copy number in sequence data.

**Table 5.** Starting values of parameters estimated by pseudo-maximum likelihood.

| Variable | Starting value | Lower bound | Upper bound |
| --- | --- | --- | --- |
| $\mu_2$ | $\bar{\mu}$ | $-\infty$ | ∞ |
| $\sigma_2^2$ | $\bar{\sigma}_2^2$ | 0 | ∞ |
| $\sigma_0^2$ | 1.5 | 0.01 | ∞ |
| $\pi_0$ | 0.01 | 0 | 0.05 |
| $\pi_1$ | 0.025 | 0 | 0.2 |
| $\pi_3$ | 0.001 | 0 | 0.2 |
| $\pi_4$ | 0.001 | 0 | 0.05 |

### Comparison of CNVRs to those identified in the literature

CNVR sequences were masked against AgResearch ovine repeat database isgcandrepbase2 and BLASTed against btau_4.0 using BLAST parameters ‐F 'm D' ‐U T-Z 2000 [69] to obtain their positions on the genome. Genomic positions on btau_4.0 of CNVs identified from seven other sheep and cattle studies [28, 21–23, 25, 19, 20] were obtained. An overlap of 1bp or more between autosomal CNVRs identified in this study and these seven other studies was used to give an indication as to how many CNVs from other studies we were able to detect and how many of the CNVs detected in this study were also reported in the other studies.

### Overlap between autosomal CNVRs and genes

CNVR sequences were masked (isgcandrepbase2) and BLASTed (parameters ‐F 'm D' ‐U T ‐Z 2000) against OARv3 to obtain their positions on the genome. Positions of the coding sequence of genes on OARv3 were provided by BGI (personal communication, Rüdiger Brauning). Overlap between autosomal CNVRs and the coding sequence of genes were determined. CNVRs that overlapped gene coding sequences by 1bp or more were used to derive the proportion of CNVRs overlapping genes. Overlap with the agouti signalling protein and adenosylhomocysteinase genes were used as a positive control, as this locus is observed as duplicated in the sheep genome [27].

A Monte Carlo simulation was set up to randomly distribute the CNVRs throughout the sheep genome and to create a distribution of the expected proportion of deletion CNVRs and duplication CNVRs overlapping genes (by at least 1bp). One hundred iterations were run to generate 100 expected proportions for both duplications and deletions. For both duplication and deletion CNVRs, the observed proportion was ranked along with the 100 simulated proportions and a two-tailed empirical p-value was calculated.

## Declarations

### List of abbreviations

Absl2r ‐: absolute log_2_ ratio
bps ‐: base pairs
BWA ‐: Burrows-Wheeler Alignment
CGH ‐: comparative genomic hybridisation
CNV ‐: Copy number variants
CNVR ‐: CNV regions
IMF ‐: International Mapping Flock
indels ‐: INsertions/DELetionS
ISGC ‐: International Sheep Genomics Consortium
Kb ‐: kilobase
Mb ‐: megabase
Oligo aCGH ‐: oligonucleotide CGH array
QTL ‐: quantitative trait loci
SD ‐: Segmental duplications
SLE ‐: systemic lupus erythematosus

### Ethics

This study was carried out in strict accordance of the guidelines of the 1999 New Zealand Animal Welfare Act and was approved by the AgResearch’s Invermay, Animal Ethics committee (applications AE154 and AE10879).

### Availability of data and materials

The UMD3_OA ovine genome build (an in-house AgResearch comparative sheep genome assembly, built using cattle reference genome UMD3 is accessible at www.sheephapmap.org/CNV/. Whole genome resequencing files are deposited in the NCBI short read archives (accessions <u>SRX150284</u>, <u>SRX150292</u>. <u>SRX150299</u>. <u>SRX150330</u>. <u>SRX150341</u>. <u>SRX150350</u>). The raw and SEGMNT processed HDaCGH data are deposited in figshare https://dx.doi.org/10.6084/nn9.figshare.3007282 along with a description file.

### Competing interests

The authors declare that they have no competing interests.

### Funding

The authors wish to acknowledge the financial contribution made by Ovita Limited and USDA grant AFRI 2009-03305 for providing funding for a large part of this work.

### Authors’ contributions

GMJ carried out the research and wrote the manuscript. MEG, MAB and JCM participated in study design, provided input on analysis and revised the manuscript. JWK provided access to data and revised the manuscript. KGD consulted on statistical analysis. BA was involved in sequence analysis. RB carried out and advised on bioinformatic processes and was involved in the design of the Roche NimbleGen 2.1M CGH array. NC revised the manuscript. All authors read and approved the final manuscript.

## Acknowledgments

The authors also wish to acknowledge the International Sheep Genomics Consortium for access to sheep samples, whole genome sequence from these samples and early access to genome assemblies.

## Additional files

**Supplementary table 1**

A description of the overlap of the platforms the various animals have been genotyped on.

**Supplementary table 2**

A list of the positions of the CNV regions on genome build UMD3_OA

**Additional file 1**

A description of the results of the cross platform (385K CGH and SNP50 chip) verification of CNV regions.

